# The gradient model of brain organization in decisions involving ‘empathy for pain’

**DOI:** 10.1101/2021.11.28.470235

**Authors:** Karin Labek, Elisa Sittenberger, Valerie Kienhöfer, Luna Rabl, Irene Messina, Matthias Schurz, Julia C. Stingl, Roberto Viviani

## Abstract

Influential models of cortical organization propose a close relationship between heteromodal association areas and highly connected hubs in the default mode network. The ‘gradient model’ of cortical organization proposes a close relationship between these areas and highly connected hubs in the default mode network, a set of cortical areas deactivated by demanding tasks. Here, we used a decision-making task and representational similarity analysis with classic ‘empathy for pain’ stimuli to probe the relationship between high-level representations of imminent pain in others and these areas. High-level representations were co-localized with task deactivations or the transitions from activations to deactivations. These loci belonged to two groups: those that loaded on the high end of the principal cortical gradient and were associated by meta-analytic decoding with the default mode network, and those that appeared to accompany functional repurposing of somatosensory cortex in the presence of visual stimuli. In contrast to the nonspecific meta-analytic decoding of these loci, low-level representations, such as those of body parts involved in pain or of pain itself, were decoded with matching topics terms. These findings suggest that task deactivations may set out cortical areas that host high-level representations. We anticipate that an increased understanding of the cortical correlates of high-level representations may improve neurobiological models of social interactions and psychopathology.

## Introduction

Early models of cortical organization based on data on axonal projections from nonhuman primates have drawn attention to a qualitative distinction concerning high-level cortical areas, such as those in prefrontal cortex and heteromodal association areas (Goldman-Rakic, 1988; Mesulam, 1990). These areas are characterized by sparse long-range connectivity, absent in upstream unimodal cortical areas dedicated to progressively encoding incoming stimuli. A related notion is that of cortical ‘converge zones’ to integrate information that is fragmentally represented in unimodal areas (Damasio, 1989). These models suggest a hierarchical organization to achieve increasing abstraction and combination of information from multiple modalities (Mesulam, 1990).

More recently, the study of connectivity in man with neuroimaging methods has delivered data largely consistent with this distinction (Yeo et al., 2011). Graphic models of connectivity, for example, have identified a network of ‘core hubs’ located upstream of sensory areas (van den Heuvel & Sporns, 2013; see also Bassett & Bullmore, 2006; Sporns, Honey, & Kötter, 2007; Braga, Sharp, Lesson, Wise, & Leech, 2013). Among studies of resting state connectivity, the ‘gradient model’ has recently characterized the cortex in terms of a collection of connectivity gradients, where each cortical point occupies a position representing topological distance to the gradient poles. The main gradient contains the highly connected core hubs of heteromodal association areas at one pole and sensory/motor areas at the other (Margulies et al., 2016; Smallwood et al., 2021), capturing the functional gamut from perception and action to high-level cognitive functions. Confirming earlier observations (Buckner et al., 2009), these studies also found the heteromodal association pole to be partially coextensive with the default mode network, a set of cortical areas deactivated by cognitively demanding tasks (Shulman et al., 1997; Raichle et al., 2001). However, it remains difficult to characterize the computations that take place in these core hubs (Avena-Koenigsberger, Misic, & Sporns, 2018).

Because of their topological distance to primary sensory and motor areas (Mesulam, 1990), the hubs at the high end of the cortical gradient may be involved in computations that require decoupling from the environment, as in the creation of an internal mental space populated by high-level semantic representations (Binder et al., 1999; Buckner, Andrews-Hanna, & Schacter, 2008; Spreng, Mar, & Kim, 2008; Smallwood et al., 2013, 2021). Recent meta-analyses also suggest these hubs to be involved in social cognition (Schurz et al., 2021), confirming observations of previous studies (Spreng, Mar, & Kim, 2008; Schurz, Radua, Aichhorn, Richlan, & Perner, 2014). A second gradient in the same decomposition places visual and sensorimotor areas at opposite ends (Margulies et al., 2016).

In the present study we employed visual stimuli used in the classic ‘empathy for pain’ neuroimaging literature (Jackson, Meltzoff, & Decety, 2005; for review, see Lamm, Decety, & Singer, 2011) to probe the cortical organization of representations of anticipated pain in others at high levels of abstraction (Figure 1A). These stimuli show hands or feet in scenes in which injuries of different types are imminent (painful images), or control scenes where the same situation does not imply a possible injury (neutral images). Our aim was to provide evidence for the embedding of these representations within the large-scale cortical organization of the brain described above, including their relationship with the default mode network probed by task deactivations, and their position within the cortical gradient defined by the gradient model. We intended to use this position to reciprocally inform the interpretation of function of cortical areas, and of the nature of computations at the high end of the cortical gradient.

**Figure 1.**
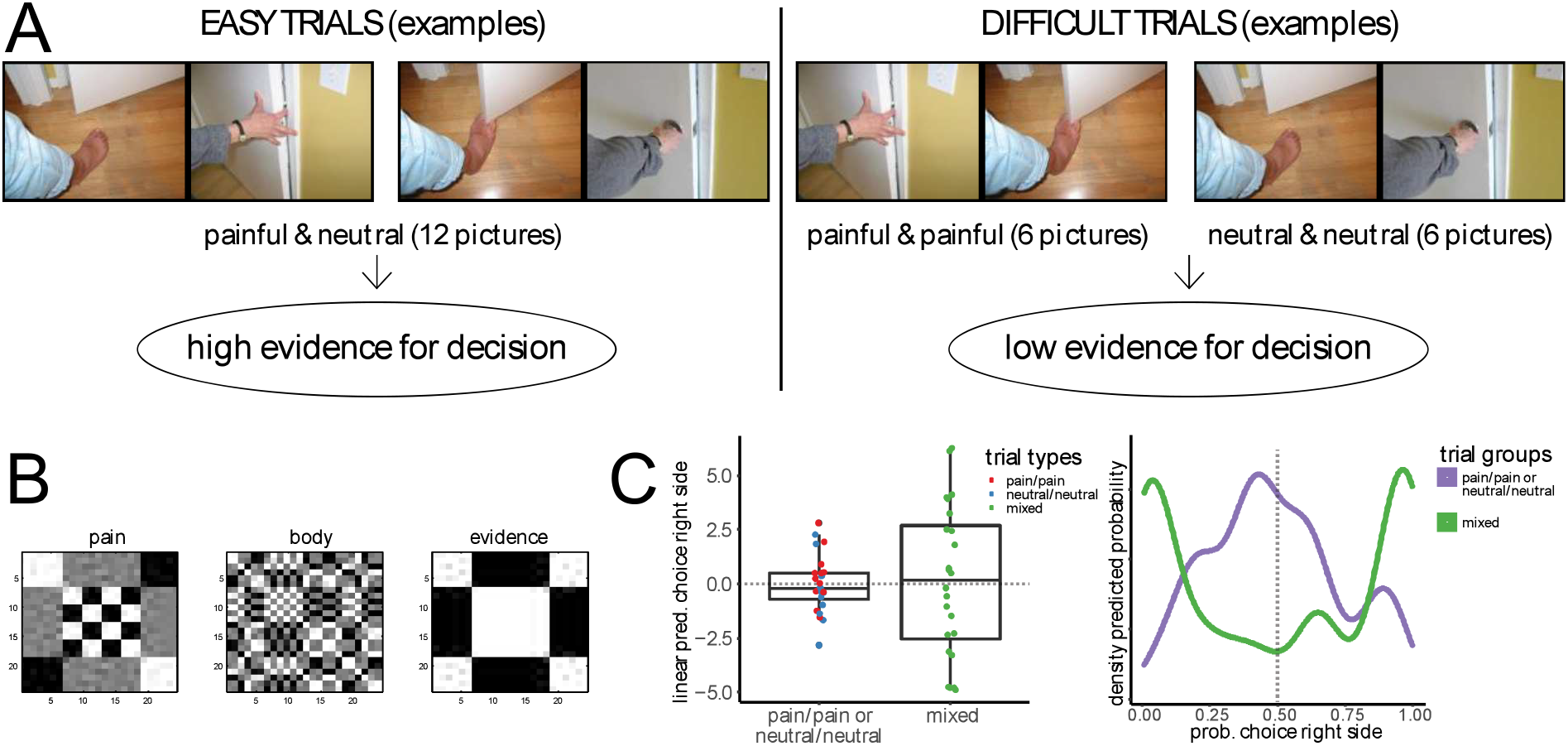
A: Examples of images from Jackson et al. (2005), showing trial arrangements leading to trials with high evidence for decisions (one image painful, one neutral) and trials with low evidence (both images of the same type). Note that this design makes it possible to select trial mages so that in the contrast for decision evidence the same images are involved in both terms of the contrast. B: representational similarity maps used in the study (rearranged to highlight structure). C: boxplot of linear predictions of right-side choice (random effects of trials) in mixed trials and painful/painful or neutral/neutral trial (left), and density of predicted probabilities of choice of right side for trials in the two conditions of the experiment (right).

Beside the widely used pain vs neutral contrast, we used two distinct approaches to characterize the cortical correlates of representations of pain. The first draws on a binary decision making design (Heekeren, Marrett, & Ungerleider, 2008; Rangel & Clithero, 2014) to identify areas associated with evidence for imminent physical pain in the visual stimuli. The decision making framework provides an explicit formulation of the high-level representations based on which decisions are made: the evidence for the decision (Heekeren et al., 2008; Shadlen & Kiani, 2013). These representations may be the furthest removed from the encoding of specific stimuli, as they may be directly translated into motor responses (Cisek, 2012), and are therefore candidates for being hosted at the high end of the cortical gradient (Viviani, Dommes, Bosch, & Labek, 2020). Furthermore, they emerge in a distributed network that integrates signals from different parts of the brain into a lowdimensional criterion (Cisek, 2012; Yoo & Hayden, 2018; Fine, Yoo, Ebitz, & Hayden, 2021). In the current study, participants were presented with two visual scenes and were asked to rate them with the question “which situation is more painful?” We assumed that evidence for decisions was large when one scene represented a painful situation and the other represented a non-painful or neutral situation (‘mixed trials’). In contrast, we reasoned that there was little evidence to decide between two images when both depicted the same type of situations (painful/painful or neutral/neutral, Figure 1A). Mixed trials and trials with the same type of situations correspond to the ‘easy’ and ‘difficult decisions’ trials of the perceptual decision making literature (Heekeren et al., 2008).

The second approach took stock of the evidence that semantic representations of stimuli are encoded in the cortex as vectors of activity in a local vector space (Haxby, Connolly, & Guntupalli, 2014), whose directions vary in each individual. To seek evidence of the pattern of activity associated with the property of stimuli we used a searchlight-based representational similarity analysis approach (RSA, Kriegeskorte, Mur, & Bandettini, 2008). This approach seeks evidence of the encoding of constructs and of semantic properties of stimuli by assessing the concordance of the representational similarity of these properties and the pattern of covariance in the brain signal elicited in each trial. We computed the representational similarity of each trial for recognizable figural elements of the images (such as feet or hands) as well as in terms of high-level features related to the final criterion for the decision (the imminent possibility of pain, i.e. the criterion to compute the evidence for a decision in the trial, Figure 1B).

Our hypotheses were designed to provide insights about the computations hosted at the high-end terminals of the cortical gradients. First, we hypothesized that the high-level representations elicited by the evidence for decisions would co-localize with task deactivations (taken as functional localizers of the DMN) and areas at the high end of the gradient model, consistently with their high degree of abstractness. We also hypothesized that encoding of simple categories, such as body parts, would be located in more upstream areas, due to their more concrete character. These two hypotheses characterize two types of computational outcomes corresponding to the different connectivity of the cortex at different points of the gradient: distributed connectivity for evidence for decisions (Viviani, Dommes, Bosch, & Labek, 2020), and local connectivity for categorization. We had no information from previous studies to form hypotheses on the RSA of the criterion for decisions (the imminent possibility of pain), but we assumed it to be located near evidence areas.

## Results

### Behavioral data

There were an average of 0.53 missed trials per participant in mixed trials and of 0.90 in pain/pain or neutral/neutral trials. This difference was significant (Poisson regression with repeated measurements, *z* = 2.13, *p* = 0.018, one-tailed). Furthermore, in mixed trials, participants gave the correct response in an average of 83.7% of trials (95% confidence interval 79.5-87.1), indicating that participants were executing the task and identifying the correct response.

Pain/pain or neutral/neutral trials had no clear correct answer, preventing the computation of an equivalent statistic. To show the difference in the existence of a clear answer in the two types of trials, we fitted a random effects logistic regression where we modelled two separate random components for trials in the mixed and in the pain/pain or neutral/neutral trials. When choices are random, the linear predictor of a logistic regression model hovers around zero, while trials with a tendency to generate a right or a left choice inflate the random component of the trial type as linear predictions for trials are correspondingly positive or negative. The variance of the random effect of mixed trial was 8.2, of pain/pain or neutral/neutral trials 2.1 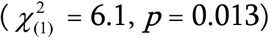, confirming that mixed trials were prevalently associated with high evidence or ‘easy decisions in contrast to trials with pain/pain or neutral/neutral images. Figure 1C shows a box plot of the linear predictions in the two types of trials, and the density of fitted trial probabilities of rightside choice. The boxplot shows the larger dispersion of linear predictions in mixed trials, as participants were more consistently choosing one of the scenes on the right or left. The density of fitted probabilities in mixed trials was bimodal, meaning that there was a tendency to select a right or left response, while the density of pain/pain or neutral/neutral trials had one mode at about 50% chance of right or left choice. However, there were a few mixed trials where participants were not entirely consistent in selecting one option, and several pain/pain or neutral/neutral trials where participants were expressing a clear preference. There was no difference in the dispersion of the random effects for pain/pain vs neutral/neural trials 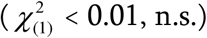.

Additional analysis of behavioural data is in the Appendix.

### Neuroimaging data

#### Evidence for decisions

In the functional imaging data analysis we first looked at the neural correlates of evidence for decisions. As in Heekeren et al. (2004), we contrasted high vs low evidence decisions, i.e. mixed trials vs trials where both scenes were of the same type. This contrast revealed a sparse network of areas involving the motor/premotor and somatosensory cortex, extending medially to the posterior cingulum (Figure 2A and Table A1 in the Appendix).

**Figure 2.**
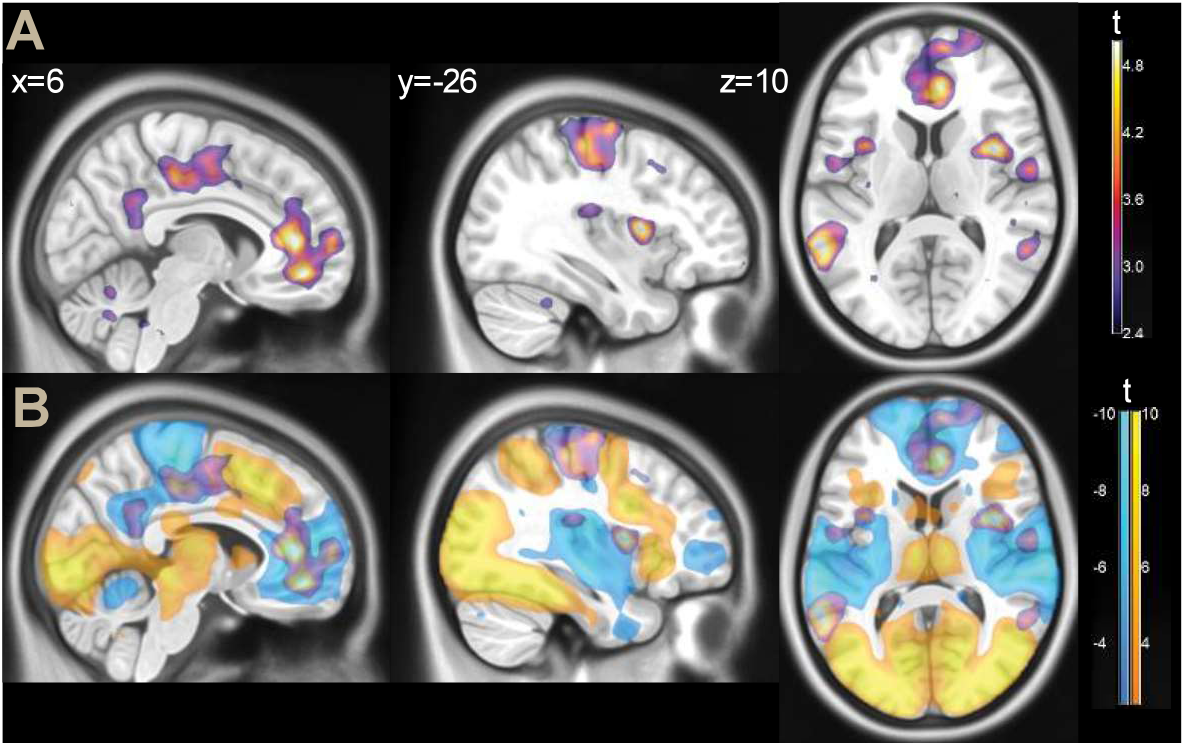
A: parametric maps, contrast high vs low evidence decision (thresholded at *p* < 0.01 for illustration purposes), overlaid on a template brain. B: the same contrast, displayed together with task activations (yellow) and deactivations (light blue, same threshold).

More anteriorly, the middle/posterior insula, and the anterior cingulum/ventromedial prefrontal cortex (vmPFC) were also involved. On the lateral aspect of the brain, the middle temporal gyrus/temporoparieal junction were also strongly associated with evidence for decisions (areas PGa and PFm in the histological classification by Caspers et al., 2006).

Several of these areas (such as the posterior cingulum and the vmPFC) are typically associated with the default mode network. We therefore looked at task deactivations, represented in light blue in Figure 2B. One can see that the effect of evidence for the decision, detected by the high vs low evidence decision contrast, involved task deactivations not only in areas classically associated with the default mode network, but also in the insula and in the motor/premotor cortex. In the insula, the association with evidence for decisions was located at the transition between the activated anterior insular and the middle and posterior portions. In the motor/premotor cortex, the same association overlapped with task deactivations, especially marked in the medial face, extending anteriorly towards the task active premotor cortex.

For completeness, we should mention that there were effects in the opposite direction (contrast high vs low evidence decisions), involving task-activated cortex in the inferior frontal gyrus and the intraparietal sulcus (see Table A2 in the Appendix). These are known effects, elicited by increases in the difficulty of cognitive tasks (e.g., Rypma, Prabhakaran, Desmond, Glover, & Gabrieli, 1999) and involved in stimulus-response mapping (Bode & Haynes, 2009; Woolgar, Thompson, Bor, & Duncan, 2011). They may represent attentional and executive processes required by the decision making task that are complementary to the activity elicited by high evidence decisions (for a discussion, see Heekeren et al., 2008). Since they are not the focus of the present study, we will not comment on them further.

#### Contrast pain vs neutral images

One question we wanted to address was the relationship of the effects of evidence for decisions with those emerging from a more traditional contrast of painful vs neutral stimuli. Because the trials with difficult decisions contained two images of the same type (painful or neutral), they could be used to estimate this contrast. The main effects of this contrast were localized in the SII/inferior parietal cortex (areas PFop, PFt, and OP1 in the classification by Caspers et al., 2006), anterior to the parietal effects of evidence and without substantial overlap (Figure 3 and Table A3 in the Appendix).

**Figure 3.**
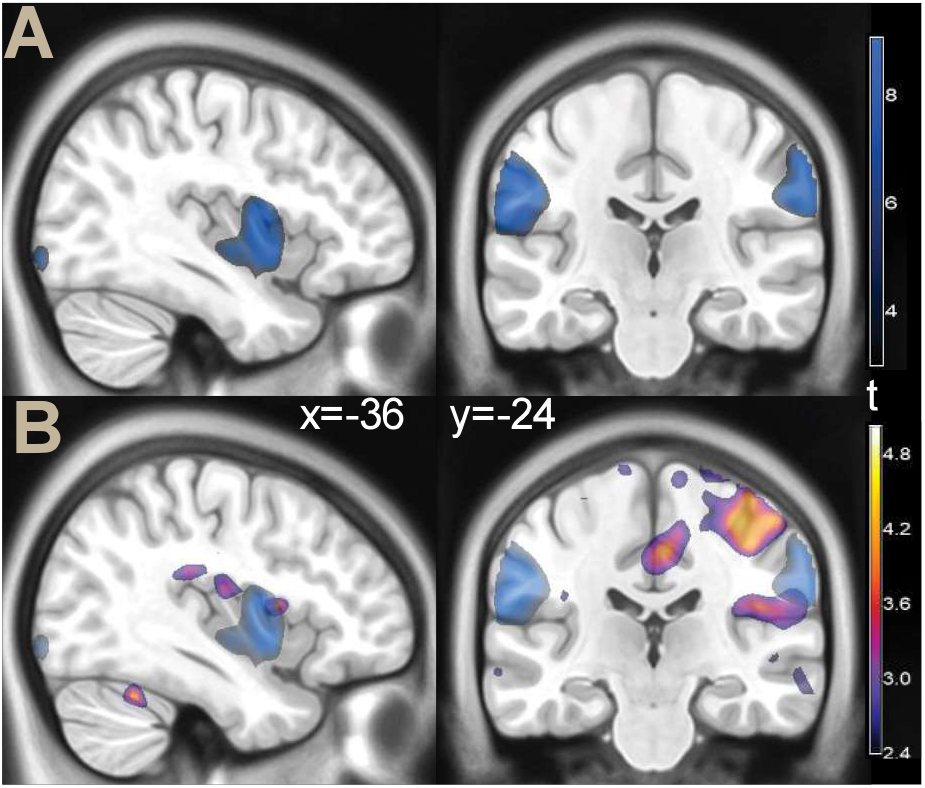
A: parametric *t* map of the contrast painful images vs neutral (illustration threshold *p* < 0.01), overlaid on a template brain. B: the same contrast (blue color), displayed together with the effect of evidence for decisions (as in Figure 2).

An additional effect of the traditional contrast was in the middle insula. While some very limited overlap with the effects of evidence was observed here, most of the activity of the traditional contrast was located ventrally to the effect of evidence. There was no relation with any of the other effects of evidence documented in Table A1.

#### Representational similarity analysis

We then turned to the investigation of representations using the RSA approach. Because the scenes of impending pain figured either hands or feet, with equal frequency across the experiment, we first computed the RSA associated with these body parts to validate the effectiveness of our approach (for more details, see Viviani, 2021). This analysis resulted in only one significant cluster of representational activity (MNI *x, y, z*: 54, -64, -2, right middle temporal gyrus, BA 37, *t* = 6.09, *p* = 0.001, peak level corrected, *p* = 0.006, cluster level corrected, 645 voxels; see Figure 4A). This cluster matched the right association area for the term ‘body’ in the automated meta-analytic tool Neurosynth (Figure 4B, Yarkoni et al., 2011; a similar map is generated by the term ‘hand’). This cluster was located at the lateral end of the large swath of task activity emanating from visual areas in the posterior occipital cortex (task activation at 54, -64, -2: *t* = 9.64, *p* < 0.001).

**Figure 4.**
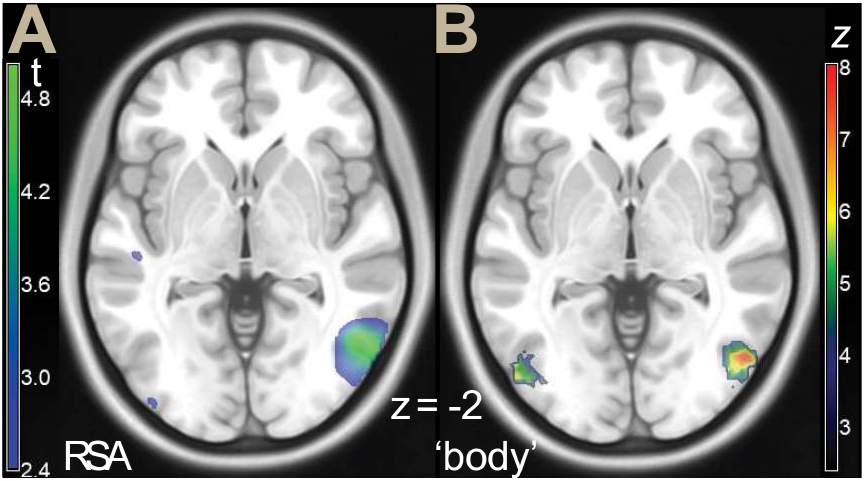
A: parametric *t* map of the RSA analysis for hands/feet (illustration threshold *p* < 0.01), overlaid on a template brain. B: Neurosynth rendering for the term ‘body’, association test FDR *p* < 0.01.

We then looked at the representation of pain, based on the distinction between impending pain and neutral images presented in the trials. This similarity map captured the presence or absence of images containing the criterion based on which choices were made. A similarity map for evidence for decisions was used as a confounder and partialled out (Figure 1B). This analysis showed extensive bilateral associations in the primary and associative sensory cortex (Figure 5A, Table A4).

**Figure 5.**
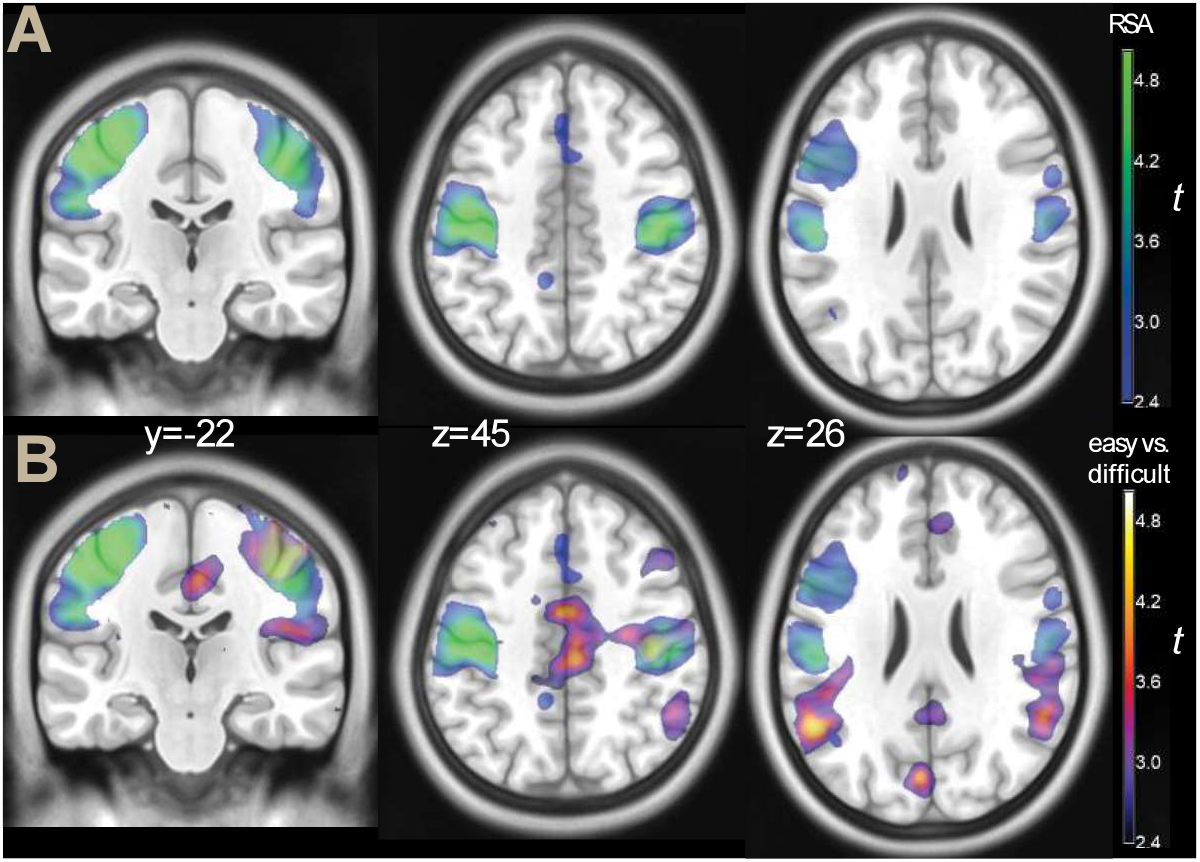
A: Parametric map of the RSA for pain (in blue-green), and B: the same map, shown with the contrast high vs low evidence decision (the same as shown in Figure 2, in blue-yellow), overlaid on a template brain. Both contrasts were thresholded at *p* < 0.001 for illustration purposes.

When shown together with the neural correlates of evidence for the decision (Figure 5B), the involvement of the somatosensory cortex in the RSA for imminent pain appeared to complement the prevalently motor involvement of evidence for decisions on the other side of the central sulcus. In the most ventral part, they reached the painful vs neutral effects of a traditional contrast in sensory association cortex, but were much more extensive than this latter.

#### Decoding analysis

To situate the areas identified in the study within the existing corpus of neuroimaging data, we looked at these four effects (evidence for decisions, contrast pain vs neural, RSA for body parts, RSA for imminent pain) with the automatic decoding tool provided by the Neurosynth website (see Methods for details). The decoding analysis, shown in Figure 6A, confirmed that the effect of evidence for decisions involved mainly areas associated in previous studies with somatosensory and motor functions, together with the related body representations, and with the default mode network. An additional set of topics terms identified by the decoding analysis involved social cognition/theory of mind. The RSA of imminent pain was substantially a subset of these terms, centered on somatosensory and motor functions. In contrast, the pain vs neutral contrast and the RSA of body parts were decoded with topics terms that were specifically related to the content of the present analyses. The pain vs neutral effect identified topics terms associated with pain, somatosensory function, and, less specifically, with negative emotion. The topics term ‘empathy’ was also elicited by this decoding, presumably in connection with ‘empathy for pain’ studies. As anticipated in Figure 4, the RSA of body parts identified topic terms associated with visual encoding and identification of body parts.

**Figure 6.**
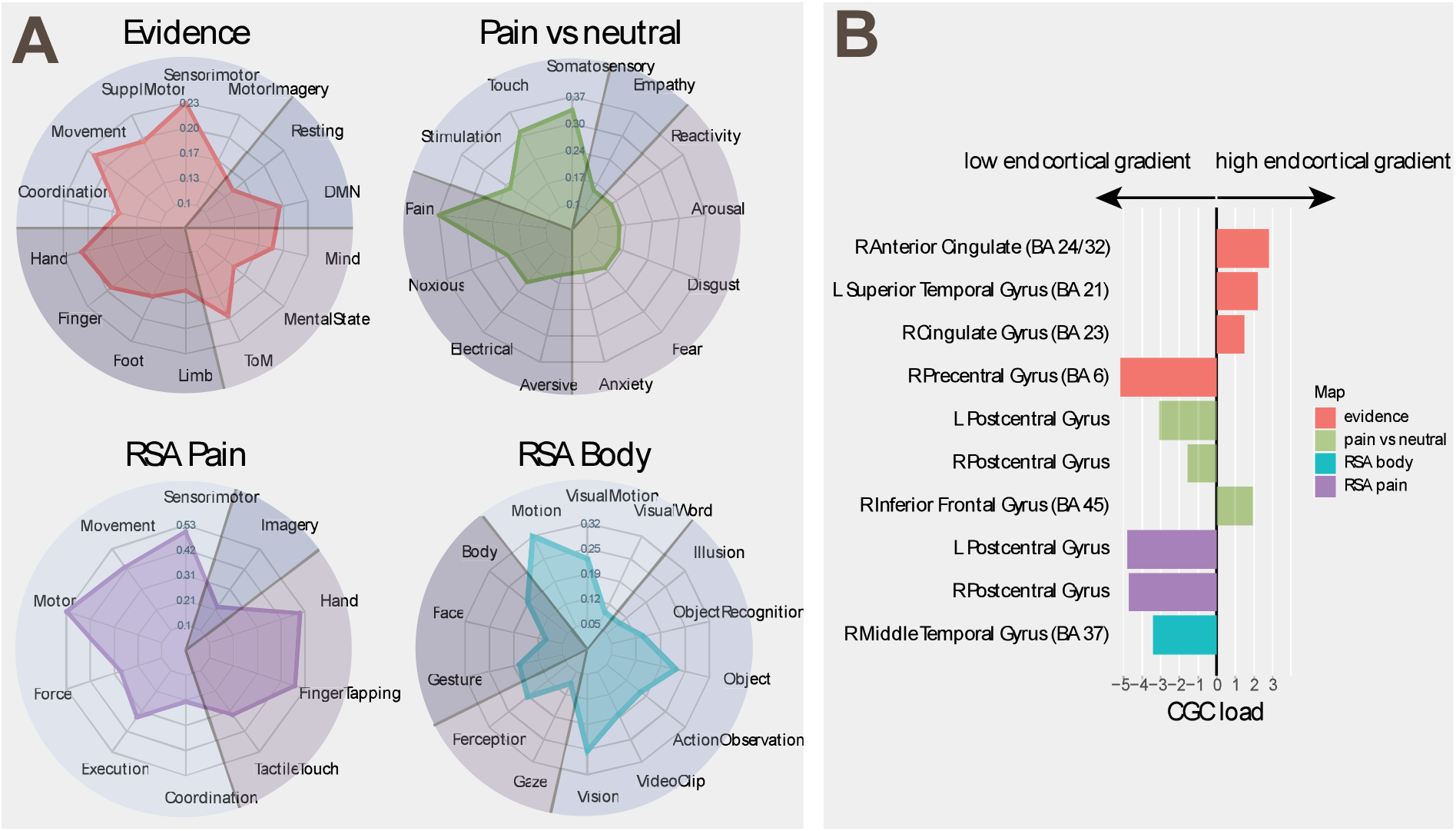
A: Decoding analysis of the effects of evidence for decisions and the contrast pain vs neutral images (top), and of RSA of imminent pain and body parts (bottom). DMN: default mode network; TOM: theory of mind. B: Average load of significant clusters on connectivity gradient component. CGC load: average load on connectivity gradient component: L, R: left, right. The anatomical labels refer to the peak of the cluster (from www.talairach.org). For more information about the clusters and their peaks, see the Tables in the Appendix.

#### Average load on connectivity gradient component

To verify the position of the areas identified in our analysis within the main gradient of cortical organization defined by connectivity (Margulies et al., 2016), we computed the mean component values of this gradient for statistically significant clusters (Figure 6B). This gradient is anchored at one end to areas serving mainly sensory and motor functions, and at the other end to heteromodal association areas serving abstract cognitive functions involving the default mode network (Margulies et al., 2016). We found that the clusters associated with evidence for decisions were characterized by high cortical gradient loadings, with the exception of the sensorimotor cluster, which loaded on the other end of the gradient. The clusters identified by the pain vs neutral contrast were also located toward the low end of this gradient. An exception was the cluster in the right inferior frontal gyrus. Finally, both RSAs gave clusters at the low end of the gradient, as appropriate for sensory representations.

## Discussion

In the present study we used a decision-making paradigm and RSA as strategies to make explicit the most abstract representations that the brain must compute to execute the task. Computationally, one may conceive of decisions as the extraction of increasingly high-level features of the stimulus up to a final stage in which neural activity computes the distance to a discriminating hyperplane, which is larger the greater the evidence for the decision (Yoo & Hayden, 2018). In the present study, areas associated with evidence for decisions were consistent with those detected in previous studies that used visual stimuli (perceptual decision making, the posterior cingulum/precuneus, the posterior temporal gyrus and adjacent parietal areas, Heekeren et al., 2004), and with the social nature of the criterion used in the decision (the posterior cingulum, the middle insula, Viviani, Dommes, Bosch, Stingl, & Beschoner, 2018). Also areas reported in studies of preference-based decision making were detected (the posterior cingulum, the vmPFC, Clithero & Rangel, 2014). In summary, recruitment of perceptual decision-making areas was consistent with the use of visual stimuli, and of those of social decision making with the criterion used in the decision.

Our first hypothesis was that these areas would involve the default mode network. Meta-analytic decoding confirmed that this was the case for some of these areas: the vmPFC and the posterior cingulum, the temporoparietal junction. These areas also loaded at the high end of the gradient model. The results of meta-analytic decoding may be viewed with caution, given that they necessarily reflect how activations were interpreted in the original studies. However, independent evidence on a specific role comes from the finding that these areas were co-located with task deactivations, or at the boundary between task activations and deactivations, replicating the findings of Viviani et al. (2020). This finding is consistent with the notion that the representations computed as evidence for deciding between options are hosted in the highly connected network located coextensively or in proximity of the default mode network, and that are identified by the cortical gradient model as convergence hubs. Furthermore, confirming our second hypothesis, the close association with task deactivations and the default mode network distinguished the areas associated with decision evidence from those elicited by a classic pain vs neutral contrast, or the RSA of body parts.

However, not all areas that were here associated with evidence for decisions followed this pattern. The sensorimotor cortex, in particular, is not part of the default mode network and is not located at the high end of the first gradient component. However, its recruitment is reported in studies of perception-based decision making that have demonstrated the emergence of a signal associated with evidence for decisions in the motor areas, among others (Gold & Shadlen, 2002; Shadlen & Kiani, 2013), leading to the inclusion of these areas in the network associated with evidence for decisions (Cisek, 2012). While sensorimotor areas are characterized by poor long-range connectivity in studies based on diffusion tractography (Hagmann et al., 2008; Gong et al., 2009), they are included among the highly connected hubs in functional connectivity studies (Achard, Salvador, Whitcher, Suckling, & Bullmore, 2006), confirming a role in the connectivity core. In the principal decomposition of connectivity data, the sensorimotor areas and the adjacent parietal operculum constitute the terminal end of the second component of the cortical gradient (Margulies et al., 2016).

Likewise, in the insula, a key area for encoding pain sensation (Lamm et al., 2011; Timmers et al., 2018; Jauniaux et al., 2019; Peyron & Fauchon, 2019; Soyman et al., 2021) but not part of the default mode network, the locus associated with evidence for decisions was located in the middle portion, half-way between the active anterior and the mildly deactivated middle/posterior portions. The task deactivation gradient in the anteriorposterior direction follows connectivity gradients reported in the insula (Cerliani et al., 2012; Royer et al., 2020; see also the third gradient component in Margulies et al., 2016). However, for both posterior insula and somatosensory cortex, the interpretation of task deactivations is made difficult by activity decrements due to the visual modality of stimuli presentation (Langner et al., 2011).

When using RSA, we found that representations of imminent pain, which were not evidence for decisions but the criterion based on which this evidence could be computed, were co-localized in the sensorimotor cortex and in the sensory association area. The decoding analysis of these areas confirmed the prevalent association with motor and sensory tasks of previous studies. There are two possible interpretations of this finding. One is that the prevalent localization of its peaks in the sensory cortex and the extension into sensory association areas is a clue to the role of these areas in the representation of empathy for pain. Considerable evidence links the sensory association area to decoding emotional expressions in others (Adolphs, Damasio, Tranel, Cooper, & Damasio, 2000; Keysers, Kaas, & Gazzola, 2010). Another interpretation is that this cortex may be representing information for the decision generically, much like the motor cortex represents information about the choice to be taken irrespective of the features that determine this choice. In favor of this latter interpretation are the results of a recent RSA analysis of a classic cognitive Stroop task, which found a similar correlate of cognitive control in this region (Freund, Bugg, & Braver, 2021), even if emotion recognition played no role in that study. Together with motor or premotor areas, the postcentral sensory cortex also appears in studies of categorization applying machine learning approaches (Li, Ostwald, Giese, & Kourtzi, 2007; Davis & Poldrack, 2014; Braunlich & Seger, 2016) or tracking distance to decision boundaries (Seger, Braunlich, Wehe, & Liu, 2015) that required discrimination between abstract patterns or food categories. Because the task inherent in these discriminations did not involve information of social-cognitive nature, the involvement of postcentral sensory cortex may be non-specific for the decision type.

In contrast, the results of the RSA of a concrete feature of the visual scenes, the displayed body parts, were much more specific, and matched the visual association area for the topics term ‘body’ in the meta-analytic decoding. It also closely replicated the findings of Jackson et al. (2005), obtained with the same stimulus material using a traditional contrast approach. This portion of the lateral occipital cortex that has been univocally associated with encoding task-relevant visual object features in conventional studies (Edelman, Grill-Spector, Kushnir, & Malach, 1998; Downing, Jiang, Shuman, & Kanwisher, 2001; Grill-Spector, Kourtzi, & Kanwisher, 2001; Braunlich, Liu, & Seger, 2017). This area was also activated by the task, in contrast to those involved in evidence for decisions. Likewise, the effects of the contrast pain vs no pain involved the anterior inferior parietal /somatoform association area, as in meta-analyses of ‘empathy for pain’ studies (Timmers et al., 2018; Jauniaux, Khatibi, Rainville, & Jackson, 2019). Hence, in interpreting function, the issue arises if the same criteria should be applied to upstream perceptual areas, which compute increasingly abstract features but are still related to the perceptual modality involved, and to downstream areas linked together by long-range connectivity and involving, according to the gradient model, the default mode network. In a computational perspective, it has been suggested that downstream areas may coordinate to provide low-dimensional priors or schematic information to disambiguate signal from upstream areas (Hinton & Zemel, 1994; Friston, 2005). What may characterize them, relative to feedback involving the extraction of features within a sensory modality (Rao & Ballard, 1999), is that the computation emerges from the interaction with other network hubs, recruited as appropriate, thus appearing in studies with heterogeneous content and defying a simple functional characterization.

Models proposed in the literature to explain recruitment of vmPFC, found here to be associated with evidence for decisions and part of the default mode network (Shulman et al., 1997), exemplify these interpretive challenges. Studies of preference-based decisionmaking tasks have proposed vmPFC to compute the integrated value and costs of a choice into a single summary criterion (Levy & Glimcher, 2012; Rangel & Hare, 2010), irrespective of whether participants made the choices for themselves or another person (Kross, Berman, Mischel, Smith, & Wager, 2011). Recent analyses of the literature, however, have proposed a more general role for vmPFC, not limited to the computations of subjective value (Rudebeck & Murray, 2014; Wilson, Takahashi, Schoenbaum, & Niv, 2014; Ciaramelli, De Luca, Monk, McCormick, & Maguire, 2019; Mack, Preston, & Love, 2020; Zhou, Gardner, & Schoenbaum, 2021), consisting in selecting the information, hosted elsewhere, that is relevant to compute predictions of the consequences of responses. Recruitment of vmPFC in the present study is consistent with these revised models, especially with views of the role of the orbitofrontal cortex in decision making that emphasize its role in computing predictions of future outcomes (Rudebeck & Murray, 2014). In the stimulus set used in the present study, the pictures suggested an injury about to happen, but no actual injury was shown. In contrast, vmPFC was not associated with evidence for decisions in a social cognition task in which all relevant information was contained in the facial expression of the displayed individuals (Viviani et al. 2018). These findings are consistent with vmPFC providing top-down disambiguating schematic information, where the disambiguation consists of the computation of a predictive criterion.

More posteriorly, the evidence for decisions was associated with activity in the temporoparietal junction and posterior temporal areas at the boundary to task deactivations. This involvement was responsible for the inclusion of theory of mind terms in the decoding analysis of this contrast (Caspers, Zilles, Laird, & Eickhoff, 2010). This finding is consistent with the model advanced by Schurz et al. (2021), according to which the cortical substrates of social cognition largely overlap with the default mode network/heteromodal association areas of the gradient model (Schilbach, Eickhoff, Rotarska-Jagiela, Fink, & Vogeley, 2008; Spreng et al., 2008). In the interpretive perspective adopted here, however, functional attributions of the temporoparietal junction would differ from those of more upstream areas such as the lateral occipital cortex, here recruited for the representation of body parts, or the somatosensory association cortex, associated in the literature with the representation of pain. Instead, its recruitment may be seen as evidence of activation of schematic or conceptual information (Murphy et al., 2018; Smallwood et al., 2021), determined dynamically in interaction with other high-level connected cortical nodes. In the right temporoparietal junction, this may be schematic spatial and visual information to identify target stimuli (Shulman, Astafiev, McAvoy, d’Avossa, & Corbetta, 2007); in the left, integration of semantic information (Lanzoni et al., 2020).

Similar considerations may apply to the posterior cingulum/retrosplenial cortex, recruited here by the evidence for decisions, and a hub of numerous long-range connections with diverse cortical regions (Hagmann et al., 2008; Leech, Braga, & Shard, 2012; van den Heuvel & Sporns, 2013). This recruitment has been interpreted by some, including ourselves, as the neural correlate of psychic pain (O’Connor, 2012; Labek et al., 2017), following consistent findings in several studies (Gündel, O’Connor, Littrell, Fort, & Lane, 2003; Kersting et al., 2009; Labek et al., 2017). However, this area is also recruited together with vmPFC by studies of subjective preference in the appetitive domain (Clithero & Rangel, 2014; Viviani et al., 2019). A more general proposal about the function of this area is attentional redirection in the presence of stimuli of behavioural significance (Pearson, Heilbronner, Barack, Hayden, & Platt, 2011; Leech & Sharp, 2014). Rather than with a simple mapping between cortical areas and emotional content, these findings are consistent with accounts that emphasize the high level of conceptual elaboration of emotional information when detected in the default mode network (Satpute & Lindquist, 2019).

In summary, we found that high-level representations related to the decision task were organized in a distributed network, as in models and data regarding decision making studies. Furthermore, they appeared to involve two types of task deactivations: one related to areas typically associated with the default mode network, as postulated by the gradient model, the other to areas that may have been repurposed to host representations of relevance for the task at hand. Accordingly, activity attributable to evidence formation appeared to involve at least two cortical gradients defined by connectivity, suggesting that our initial hypothesis, involving the principal gradient only, was too simplistic. In agreement with recent proposals (Margulies et al., 2016; Smallwood et al., 2021), we argued that the functional characterization of these areas in terms of simple mappings is unlikely, especially considering proposals in the literature on the abstract nature of high-level representations and the distributed character of their computations. This contrasts with the more univocal interpretation of areas representing more elementary categories, such as body parts, or of areas identified by the subtraction of signal in neutral from pain images.

Future studies may be able to characterize the role of the connected core network of distributed hubs in clinical models. This issue arises from the observation that personality disorders, in particular, may be associated with changes in empathic processes (Ripoll, Snyder, Steele, & Siever, 2013; Luyten & Fonagy, 2015; Sosic-Vasic et al., 2019) and poor representations of interpersonal events (Skodol et al., 2011). Furthermore, it has been noted that areas located at or in proximity of task deactivations are often recruited in tasks of emotion regulation (Viviani, 2013), suggesting a role in controlling the encoding of representations of social and emotional relevance (Viviani, 2014; Messina, Sambin, Beschoner, & Viviani, 2016). Hence, insights from connectivity studies may facilitate the integration of models of semantic processing with the clinical neuroscience literature.

## Methods

### Participants

The fMRI study was conducted at the Psychiatry and Psychotherapy Clinic of the University of Ulm, Germany, after approval by the Ethical Review Board. Healthy participants (*N* = 50) were recruited through local announcements and admitted to the study after providing written informed consent. One participant was excluded due to failure to record responses during the scan, leaving *N* = 49 for the final sample (mean age 24.7, standard deviation 5.1, range 19-47; 26 women).

### Experimental design

Participants were shown two pictures of right hands or feet in situations of anticipated physical pain or neutral outcomes (source: Jackson et al., 2005). In each trial participants were instructed to decide which situation in the two pictures could lead to the most painful outcome. We selected images from the original study showing hands or feet with equal frequency in each outcome and matching the type of injury causing pain (Figure 1A). Pictures were displayed side by side for 2.5 sec. After this time, blue dots appeared under the pictures to indicate participants could communicate their decision by pressing the left or right button, depending on the side of the anticipated painful outcome. If after 1.5 sec no button was pressed, the trial was declared as a miss. Trial onsets were generated with a Poisson interval schedule of mean intensity of a trial every 14.75 sec, bounded to a minimal interarrival time of 13 sec.

To investigate the neural correlates of evidence for decisions in an interpersonal context, we looked for a signal associated with the evidence for a decision, as in sensory-based decision making studies (Heekeren, Marrett, Bandettini, & Ungerleider, 2004). To form trials where evidence for decisions was high, we paired painful with neutral pictures as decision options. We paired painful with painful or neutral with neutral images to form low evidence trials. There were all combinations of outcomes (12 trials painful/neutral, 6 trials painful/painful, 6 trials neutral/neutral). Lateralization of pictures showing painful outcomes was balanced across the painful/neutral trials. The contrast of interest, representing high vs low evidence, subtracted the activity of painful/painful and neutral/neutral trials from the painful/neutral trials. The allocation of pictures to these combinations was swapped between participants such that in the overall sample the same pictures occurred in the painful/neutral and in the painful/painful or neutral/neutral combinations. Because the contrast high evidence vs low evidence contained the same images in the two conditions that form this contrast, it was not confounded by intrinsic properties of the images, such as their salience, the presence of imminent pain itself, or by any other property that was not matched between painful and neutral images.

The decision making design is validated by numerous functional imaging studies (for review, see Heekeren et al., 2008). The time course of neural activity in decision making tasks is well characterized by sequential sampling models, in which evidence for the available options is accumulated up to a boundary indicating sufficient evidence for one option (Gold & Shadlen, 2002; Heekeren et al., 2008; Shadlen & Kiani, 2013; for neuroimaging studies, see Ploran et al., 2007; Lim, O’Doherty, & Rangel, 2011; Wheeler et al., 2015). Here, we only focus on the existence of a neuroimaging signal correlated with the evidence for decisions. In functional imaging studies of perceptual decision making, the contrast used to detect neural activity associated with evidence for decisions is referred to as the “easy vs difficult” decisions contrast (Figure 1A), because when the evidence for decisions is large the decision is easy (Heekeren et al., 2004, 2008).

Data were collected in a Prisma 3T scanner (Siemens, Erlangen) using a T2*-sensitive echo-planar imaging sequence (TR/TE 1970/36 msec, bandwidth 1776 Hz/pixel, GRAPPA acceleration factor 2, 32 transversal slices acquired in ascending order, slice thickness 2.5 mm with a slice gap of 0.625 mm, field of view 192 with matrix size 64, giving in-planar voxel size 3×3 mm). In each participant, 182 scans were acquired after reaching equilibration giving a total scan duration of 6 min.

### Statistical analysis of behavioural data

Behavioral data were analyzed with the freely available statistical software R (http://www.R-project.org/) using the package lme4 (version 1.1-26, function *glmer*, Bates, Mächler, Bolker, & Walker, 2015). To demonstrate the tendency to consistently choose one of the options, a logistic regression model where choice of the right side button was arbitrarily coded as ‘success’ was fitted with an intercept and trials (identified by the combination of pictures of the trial) as random effects. The variance of the random effect indicates the tendency to depart from equal probability choices for right or left. To compute significance of the increased variance of trials with painful/neutral images (indicating a tendency to consistently choose right or left in high evidence trials), relative to trials with painful/painful or neutral/neutral images (low evidence trials), a likelihood ratio test was computed between a model with one random effect for all trials and a model with heteroscedastic random effects for high and low evidence trials. The density in Figure 1C was computed on random effects of fitted choice probabilities with the package ggplot2 (Wickham, 2009) using a Gaussian kernel with bandwidth 0.08.

### Statistical analysis of neuroimaging data

Functional imaging data were preprocessed with the freely available software SPM12 (www.fil.ion.ucl.ac.uk/spm) running on Matlab (The MathWorks, Natick, MA). In the main analysis, brain data were realigned, registered to a MNI template (2 mm isotropic resampling size), and smoothed (FWHM 8 mm). At the first level, the brain signal was modelled by convolving a box-car function corresponding to the trials (2.5 sec) with a canonical BOLD function. Trials of three types were coded as separate regressors: trials with a painful and a neutral scene, trials with both painful scenes, and trials with both neutral scenes. Further confounder regressors were six head movement displacements estimated from realignment, and four regressors for denoising (Behzadi, Restom, Liau, & Liu, 2007). Denoising is indicated in RSA (described below) as it may improve detection power (Charest, Kriegeskorte, & Kay, 2018). The denoising confounder regressors were obtained from the first four principal components of the data in the voxels classified as belonging to the lateral ventricles, white matter, and bone in the segmentations conducted by SPM12 as part of the registration procedure. The number of denoising regressors was based on the results reported by Kay, Rokem, Winawer, Dougherty, & Wand (2013). Data and regressors were high-pass filtered (128 sec). Serial dependency of observations was modelled with an AR(1) autoregressive term fitted to pooled residuals. Two contrasts were computed: trials with painful and neutral scenes vs trials with scenes of the same type (evidence for decisions) and trials with only painful vs trials with only neutral scenes (standard contrast pain vs control). After estimating the model in each voxel separately at the first level using the SPM12 software, contrast volumes from each subject were brought to the second level to account for the random effect of subjects. Inference was obtained using a permutation method (8000 resamples) to correct for multiple testing at the voxel and cluster level (cluster definition threshold, *p* < 0.001).

The RSA was conducted on non-smoothed data in volumes in the original space (i.e., non-registered) from coefficient estimates of a model in which each trial was modelled as a separate regressor (Kriegeskorte et al., 2008), with the same confounding covariates and autoregressive term as in the main model. We then formed the correlation matrices of the estimated model coefficients in a searchlight sphere (diameter 8 mm, 256 voxels) prior to computing the partial correlation of the off-diagonal terms of this matrix with those of the similarity matrices characterizing the trial according to the properties of the stimuli, which were computed by counting the shared properties of elements pairwise. Here, the elements consisted of the whole display of the individual trials, giving similarity matrices for trials. The properties of the trials that were used in the RSA were the anticipation of pain or no anticipation for each of the shown pictures, the body parts in the pictures (hands or feet) and the evidence for the decision (high or low evidence decisions, as a confounder; see Figure 1B). The searchlight sphere was computed with the appropriate function in the SPM12 software (spm_searchlight). We used a partial correlation approach to adjust for possible confounders (high or low evidence for decisions) in the RSA for body parts and imminent pain and to counter possible bias due to the non-orthogonality of the BOLD-convolved regressors in the design matrix (for details, see Viviani, 2021). Correlation values from the RSA were then registered with the parameters computed for the main analysis (resampling voxel size, 2 mm isotropic) and brought to the second level, where they were smoothed (FWHM 4 mm) and processed in one-sample *t*-tests using permutation (8000 resamples) to correct for multiple tests at the voxel and cluster level (cluster definition threshold, *p* < 0.001).

Coordinates in text, figures and tables of the main and the representation similarity analyses are expressed in the Montreal Neurological Institute (MNI) standard space. Overlays of statistical parametric maps were obtained with the software package MRIcroN (Chris Rorden, freely available at https://people.cas.sc.edu/rorden/mricron/index.html).

### Decoding analysis

The decoding analysis was obtained with the online tool available at the Neurosynth site (https://www.neurosynth.org, Yarkoni, Poldrack, Nichols, Van Essen, & Wager, 2011). The topics terms shown in the Results were obtained from the first 50 associations generated by the decoder after removing all anatomical terms and combining semantic duplicates (for example, ‘tactile’ and ‘touch’ were considered equivalent).

The average load on the principal gradient of cortical organization defined by connectivity was based on the data of Margulies et al. (2016), publicly available at the NeuroVault.org site (Gorgolewski et al., 2015) at the web site identifier https://identifiers.org/neurovault.image:31997. The significant clusters of the effect of evidence, of the pain vs neutral contrast, and of the RSAs of body parts and imminent pain were resampled on the space of the gradient data. Because of their intrinsically lower spatial resolution and/or smoothing, these clusters overstep the boundaries of the cortical mantle. To compute the average load on the main connectivity gradient, they were masked for the voxels where the gradient is present in the principal gradient data.

## Data availability

Parametric maps of the results of this study are available at the NeuroVault site (web identifier https://identifiers.org/neurovault.collection:11616).

## Software availability

The SPM12 add-on software used to compute the RSA using the partial correlation approach is available in a public repository (https://github.com/roberto-viviani/rsa-rsm.git).

## Acknowledgments

We would like to thank Jean Decety for kindly providing the images of impending pain we used in the experiment. We would also like to thank Sabine Wehner, Jeff Maerz, Stefanie Bernardin, and Eun-Jin Sim for technical support. The authors declare no conflict of interest.

This work was funded by collaborative grants from the Federal Institute for Drugs and Medical Devices (BfArM, Bonn, Germany, Grant No. V-17568/68502/2017-2020), by the Austrian Science Fund (FWF), Grant No. I 5903-B (ERA-PERMED Project ArtiPro), by the “Forschungsförderungsmitteln der Nachwuchsförderung 2018 der Universität Innsbruck”, project “Analyse neuronaler Populationscodes physischer und psychischer Schmerzen mit multivariaten Musterinformationsanalysen - Anwendung der ‘Functional MRI Representational Similarity Analyses’ bei funktioneller Bildgebung”, and by the Förderkreis of the University of Innsbruck (Förderkreis 1669).

## Appendix

## Neuroimaging data, contrasts

**Table A1.**
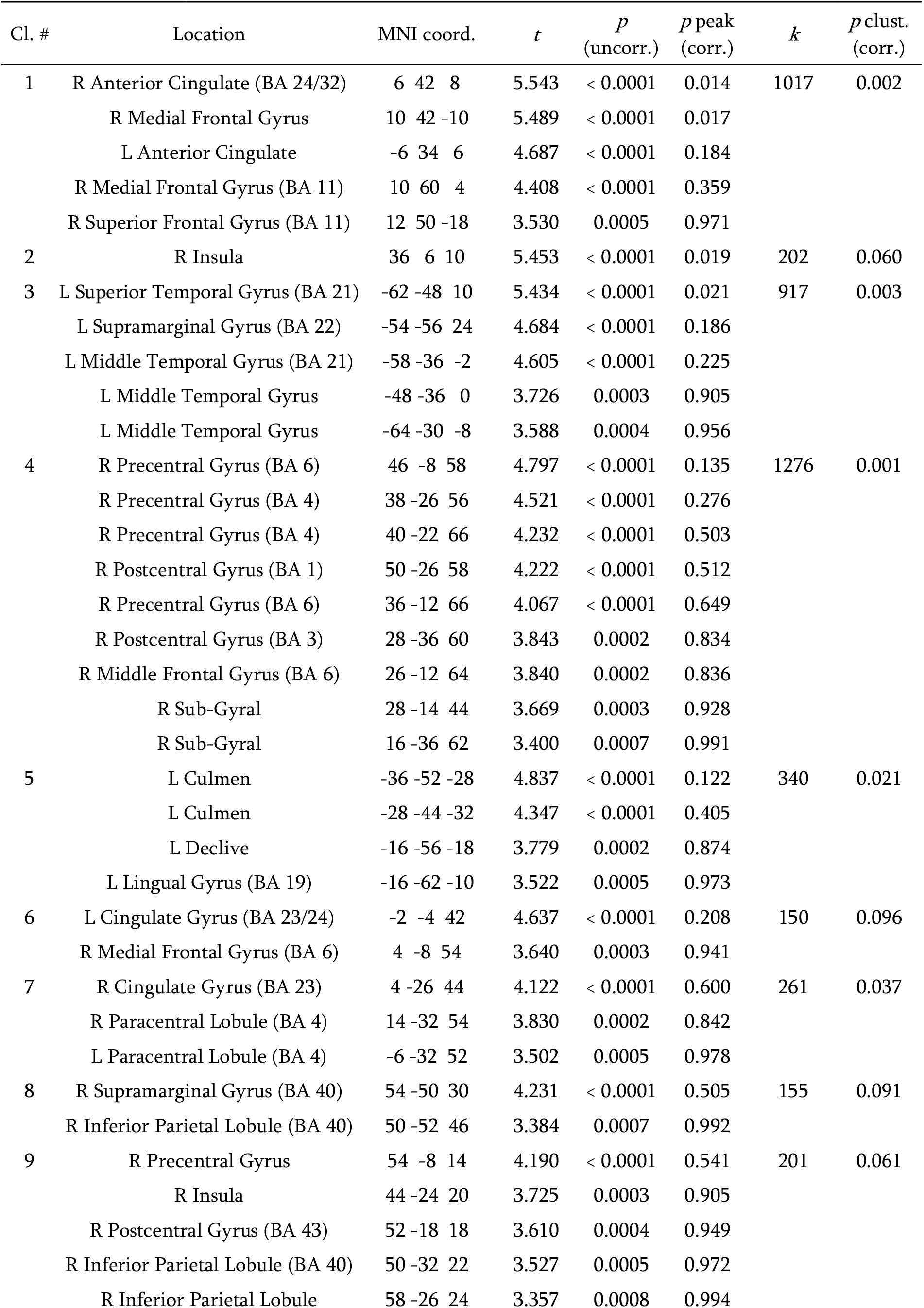
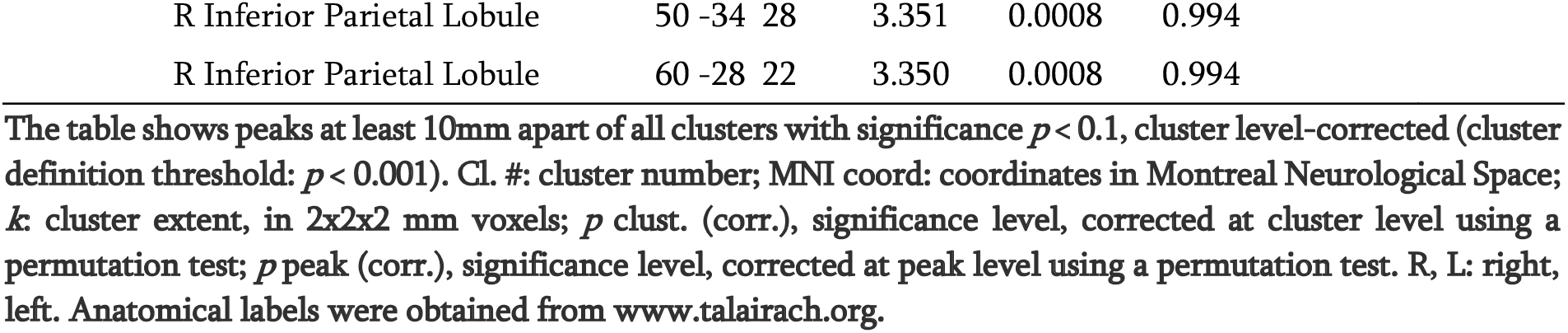
Contrast high vs low evidence decisions

**Table A2.** Contrast low vs high evidence decisions

| Cl. # | Location | MNI coord. | $t$ | $p$<br>(uncorr.) | $p$ peak<br>(corr.) | $k$ | $p$ clust.<br>(corr.) |
| --- | --- | --- | --- | --- | --- | --- | --- |
| 9 | L Inf. Frontal/ Sub-Gyral (BA 45) | -40 28 18 | 5.721 | < 0.0001 | 0.009 | 356 | 0.018 |
| 16 | L Precuneus (BA 7) | -20 -70 48 | 4.968 | < 0.0001 | 0.089 | 457 | 0.012 |
|  | L Precuneus (BA 7) | -24 -64 38 | 4.313 | < 0.0001 | 0.424 |  |  |
|  | L Superior Parietal Lobule (BA 7) | -12 -68 66 | 4.156 | < 0.0001 | 0.557 |  |  |
| 18 | L Inferior Parietal Lobule (BA 40) | -36 -46 50 | 4.573 | < 0.0001 | 0.240 | 166 | 0.076 |
|  | L Sub-Gyral (BA 40) | -34 -42 40 | 3.730 | 0.0003 | 0.893 |  |  |

The table shows peaks at least 10mm apart of all clusters with significance $p < 0.1$ , cluster level-corrected (cluster definition threshold: $p < 0.001$ ). Cl. #: cluster number; MNI coord: coordinates in Montreal Neurological Space; $k$ : cluster extent, in $2 \times 2 \times 2$ mm voxels; $p$ clust. (corr.), significance level, corrected at cluster level using a permutation test; $p$ peak (corr.), significance level, corrected at peak level using a permutation test. R, L: right, left; Inf: inferior. Anatomical labels were obtained from www.talairach.org.

**Table A3.** Contrast painful vs neutral images within difficult decisions

| Cl. # | Location | MNI coord. | *t* | *p*<br>(uncorr.) | *p* peak<br>(corr.) | *k* | *p* clust.<br>(corr.) |
| --- | --- | --- | --- | --- | --- | --- | --- |
| 1 | L Postcentral Gyrus | -60 -24 24 | 8.588 | < 0.0001 | <0.001 | 2648 | <0.001 |
|  | L Insula | -38 4 -4 | 6.735 | < 0.0001 | <0.001 |  |  |
|  | L Insula | -38 0 12 | 6.169 | < 0.0001 | 0.002 |  |  |
|  | L Inferior Frontal Gyrus (BA 6/24) | -54 6 22 | 5.796 | < 0.0001 | 0.007 |  |  |
|  | L Postcentral Gyrus | -52 -34 56 | 5.245 | < 0.0001 | 0.041 |  |  |
|  | L Insula | -38 -12 -4 | 4.682 | < 0.0001 | 0.192 |  |  |
| 2 | R Postcentral Gyrus (BA 2) | 66 -22 28 | 6.396 | < 0.0001 | 0.002 | 813 | 0.003 |
|  | R Postcentral Gyrus | 62 -24 44 | 5.738 | < 0.0001 | 0.009 |  |  |
|  | R Inferior Parietal Lobule | 60 -34 48 | 4.933 | < 0.0001 | 0.096 |  |  |
| 3 | R Inferior Frontal Gyrus (BA 45) | 48 46 4 | 6.703 | < 0.0001 | <0.001 | 377 | 0.021 |
|  | R Middle Frontal Gyrus (BA 10) | 42 60 2 | 4.047 | < 0.0001 | 0.669 |  |  |
| 4 | R Sub-Gyrus | 40 -12 -10 | 5.425 | < 0.0001 | 0.023 | 226 | 0.056 |
|  | R Parahippocampal Gyrus (BA 34) | 24 -4 -16 | 4.514 | < 0.0001 | 0.283 |  |  |
|  | R Sub-Gyrus | 38 -4 -10 | 4.392 | < 0.0001 | 0.369 |  |  |
|  | R Putamen | 30 -12 -8 | 3.804 | 0.0002 | 0.864 |  |  |
| 5 | L Extra-Nuclear | -22 -6 -12 | 4.996 | < 0.0001 | 0.080 | 170 | 0.087 |
|  | L Sub-Gyrus | -28 -14 -12 | 3.532 | 0.0005 | 0.971 |  |  |
|  | L Parahippocampal Gyrus (BA 34) | -30 4 -18 | 3.393 | 0.0007 | 0.992 |  |  |

The table shows peaks at least 10mm apart of all clusters with significance $p < 0.1$ , cluster level-corrected (cluster definition threshold: $p < 0.001$ ). Cl. #: cluster number; MNI coord: coordinates in Montreal Neurological Space; *k*: cluster extent, in 2x2x2 mm voxels; *p* clust. (corr.), significance level, corrected at cluster level using a permutation test; *p* peak (corr.), significance level, corrected at peak level using a permutation test. R, L: right, left. Anatomical labels were obtained from www.talairach.org.

## Neuroimaging data, representational similarity analysis

**Table A4.**
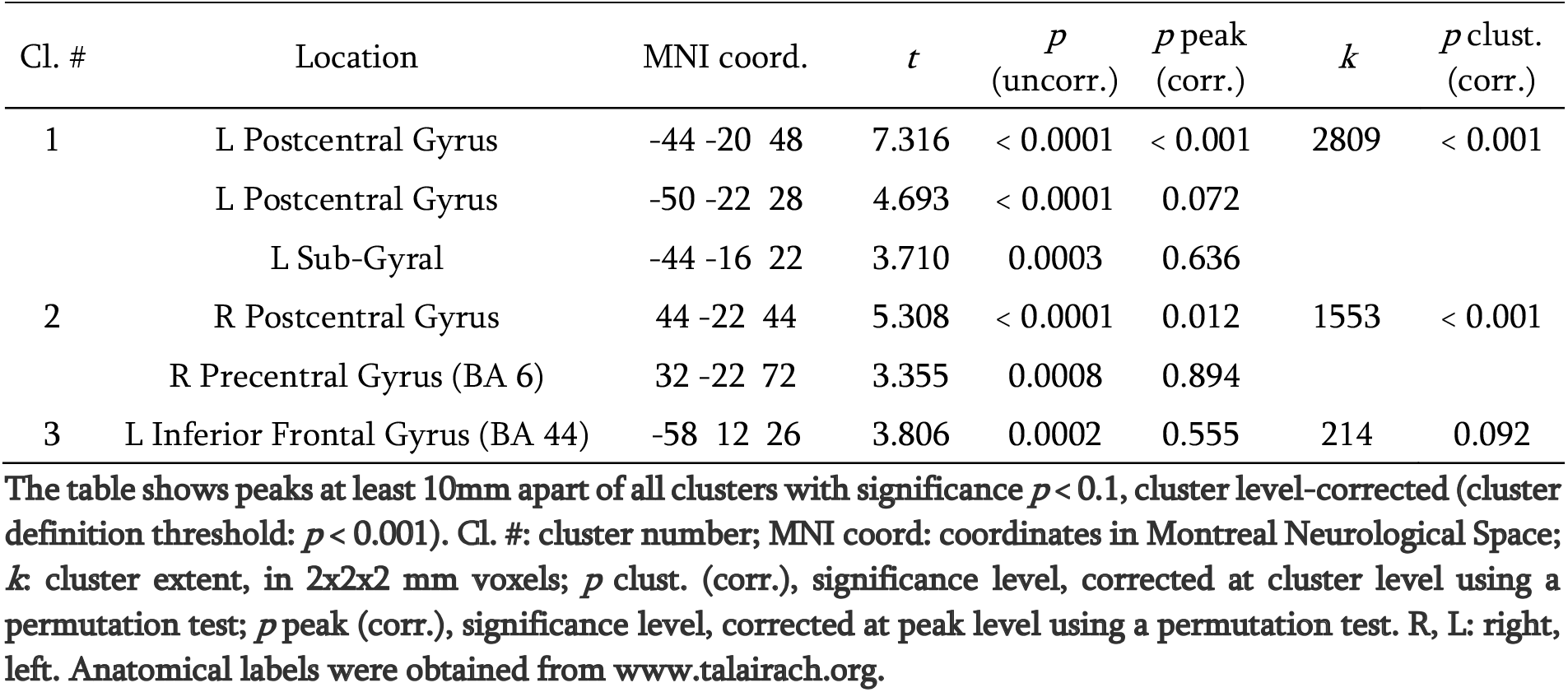
RSA analysis category ‘pain’

## Average connectivity gradient component load

**Table A5.** Average connectivity component load of significant clusters

| Map | Cl. # | Location | Peak MNI coord. | CGC |
| --- | --- | --- | --- | --- |
| high vs low evidence | 1 | R Anterior Cingulate (BA 24/32) | 6 42 8 | 2.827 |
|  | 3 | L Superior Temporal Gyrus (BA 21) | -62 -47 10 | 2.224 |
|  | 4 | R Precentral Gyrus (BA 6) | 46 -8 58 | -5.187 |
|  | 7 | R Cingulate Gyrus (BA 23) | 4 -26 44 | 1.484 |
| painful vs neutral | 1 | L Postcentral Gyrus | -60 -24 24 | -3.068 |
|  | 2 | R Postcentral Gyrus | 66 -22 28 | -1.569 |
|  | 3 | R Inferior Frontal Gyrus (BA 45) | 48 46 4 | 1.930 |
| RSA pain | 1 | L Postcentral Gyrus | -44 -20 48 | -4.787 |
|  | 2 | R Postcentral Gyrus | 44 -22 44 | -4.728 |
| RSA body | 1 | R Middle Temporal Gyrus (BA 37) | 54 -64 -2 | -3.415 |

Map: contrast or RSA analysis from which the map was obtained; Cl. #: cluster number; MNI coord.: coordinates in Montreal Neurological Space; CGC: average connectivity gradient component load. R, L: right, left. Anatomical labels were obtained from www.talairach.org. Connectivity gradient component data from Margulies et al. (2016).

## Supplementary analysis

Participants may have adopted an alternative responding strategy than the one they were instructed. It is possible to achieve perfect accuracy in easy trials by monitoring only images on one side. For example, they may have pressed the button for the left image if it is a painful image, and chosen the right image otherwise. This strategy leads to correct choices in easy trials. We show here that the pattern of responses was not compatible with adopting this strategy because of its consequences on choices made in difficult trials.

While implementing this strategy, participants would always choose the right or the left side in painful/painful or neutral/neutral trials, depending on the side they look at. If a participant follows this strategy by looking at the left image, then she would report left in all painful/painful trials and right in all neutral/neutral trials. If she looks at the right image, she would report right in all painful/painful trials and left in the neutral/neutral trials. That is, choices may have occurred equally frequently left or right in the difficult trials sets across subjects (as shown in Figure 1 of the main text), but these choices would be anticorrelated within subjects. Therefore, evidence for this strategy may be tested by looking at the correlation between counts of choices for one side (it does not matter if left or right) between the two trial sets within difficult trials (testing for a negative correlation). However, the chosen side was positively correlated between pain/pain and neural/neutral trials (Pearson correlation *r* = 0.47, *p* < 0.01), suggesting that participants tended to pick up the same side when there was too little information in favour of a specific option.

This test looks at average decisions but does not look at the possibility that individual participants may follow the alternative strategy. There were 12 difficult trials in the experiment, 6 in the painful/painful trial set and 6 in the neutral/neutral. A participants looking at the left image would score 6 left choices in the painful/painful trials and 0 in the neutral/neutral trials. If she looks at the right image, she would score 0 and 6 left choices instead. Hence, the absolute difference between left choices in these trial sets gives the tendency the participant had to follow this strategy. A participant following this strategy would score 6 at this statistic (for perfect anticorrelation), but this may be less if they realized they could do this only at some point.

In the data, the maximal absolute difference was 4, in one participant.

To see the extent to which this may be indicative of anticorrelation in this participant, we simulated experiments of 49 participants with two sets of 6 trials, and computed the number of participants who scored 4 or larger in the absolute difference of left choices between the two sets, with the left choices made independently in the two sets. In the simulation, we set the rate of left choices to the rate in the data, which was 46% (logistic regression, *p* = 0.34). We then computed the proportion of 10.000 repetitions of the simulations that gave one or more subjects with a maximal absolute difference of 4 or more. This gave 0.85, i.e., *p*(one subject or more) = 0.85. Indeed, more than half of these simulated experiments under independent choices (56%) produced more individuals with the same tendency for anticorrelated choices than in our real sample.

We then repeated the simulation with a different threshold. There were 3 participants in our data where the absolute difference was 3 or more (one of these was the participant with the difference of 4), which might have been those that adopted the strategy half-way. The simulation gave *p*(3 subjects or more) = 0.98.

This means that there was no evidence that any participant was following the alternative strategy proposed by the reviewer. On the contrary, the number of participants making anticorrelated choices was too low even relative to what we would expect from independent choices in the two trial sets, consistently with the previous finding of a general tendency for correlated choices.

